# Cell cycle arrest explained the observed bulk 3D genomic alterations in response to long term heat shock for mammalian cells

**DOI:** 10.1101/2021.12.07.471582

**Authors:** Bingxiang Xu, Xiaomeng Gao, Xiaoli Li, Yan Jia, Feifei Li, Zhihua Zhang

**Affiliations:** Beijing Institute of Genomics, Chinese Academy of Sciences, and China National Center for Bioinformation No.1 Beichen West Road, Chaoyang District, Beijing 100101, China; School of Life Science, University of Chinese Academy of Sciences, Beijing, China; School of Kinesiology, Shanghai University of Sport, Shanghai, China; Department of Cell Biology and Genetics, Core Facility of Developmental Biology, Chongqing Medical University, Chongqing 400016, China; Division of Cell, Developmental and Integrative Biology, School of Medicine, South China University of Technology, Guangzhou 510006, China

**Keywords:** Heat shock, Hi-C, chromosomal conformation, cell cycle

## Abstract

Heat shock is a common environmental stress, while the response of the nucleus to it remains controversial in mammalian cells. Acute reaction and chronic adaptation to environmental stress may have distinct internal rewiring in the gene regulation networks. However, this difference remains largely unexplored. Here, we report that chromatin conformation and chromatin accessibility respond differently in short- and long-term heat shock in human K562 cells. Interestingly, we found that chromatin conformation in K562 cells was largely stable in response to short heat shock, while showed clear and characteristic changes after long-term heat treatment with little alteration in chromatin accessibility during the whole process. We further showed *in silico* and experimental evidence strongly suggesting that changes in chromatin conformation may largely stem from an accumulation of cells in the M stage of cell cycle in response to heat shock. Our results represent a paradigm shift away from the controversial view of chromatin response to heat shock and emphasize the necessity of cell cycle analysis while when interpreting bulk Hi-C data.

## Introduction

Environmental fluctuations are part and parcel of living organisms. For example, temperature change is a common environmental stress, and the mechanism for heat shock (HS) response reported in the literature showed certain consensus patterns from yeasts to humans [1]. More specifically, cells globally repress their transcription [2, 3], RNA processing [4, 5], and translation for most genes during HS [6, 7], while many heat-shock proteins (HSP), acting as molecular chaperons, are induced, such as HSP70 and HSP90 [8, 9]. As HSPs are also prominent in various tumor types [10], HS is not just a simple model system for gene regulation studies, but also a bellwether for related diseases.

Cells may have distinct reactions, depending on the length of heat treatment [11]. This is plausible because cells must respond to both immediate environmental crises, as well as adapt to the resultant altered environment, or return to pre-stimulus status after a sufficiently long period of experiencing persistent change [12, 13]. However, to the best of our knowledge, most mechanistic studies concerning gene transcription regulation in animal cells are conducted under short-term HS [3].

Gene transcription is repressed during HS [14-17], while the behavior of chromatin spatial structures associated with gene regulation [18] in response to HS remains controversial. On the one hand, changes in chromatin spatial structure, such as genome compartments [19], topological associated domains (TAD) [20], and chromatin loops [21], have been reported to occur in association with transcriptional responses to HS in many species. For example, in yeast, upregulated HSP genes undergo a dynamic alteration of their 3D genome structure [22, 23]. In *Drosophila*, HS induces relocalization of architectural proteins from the TAD borders to the interior of TADs, causing a dramatic rearrangement in the 3D organization of chromatin [24]. In mammalians, the picture is less clear. It has been shown that chromatin loops undergo pronounced changes in human ES cells in response to HS and that the frequency of loop interactions is correlated with the level of nascent transcription [25]. On the other hand, in the human myelogenous leukemia cell line (K562), chromosomal architecture remained largely stable after short-term HS [26]. These divergent findings call for more intensive study of chromosomal architecture in reaction to HS.

At the cellular level, HS treatment can probably arrest cells at various phases of the cell cycle, depending on cell type [11, 27]. For example, in human glioblastoma cells, heat-induced expression of Hsp70 correlates with the induction of p21 and subsequent G1 cell cycle arrest [28]. In K562 cells, heat treatment caused significant accumulation of cells in the G2/M phase at 12-30h post-exposure [29].

However, the responses towards long-term HS at the levels of chromatin structure and cell cycle appear to be an underexplored topic in the literature, and it is far from clear if cells respond to short- and long-term HS in a similar manner. To address these questions, we profiled chromatin conformation and chromatin accessibility in human K562 cells under normal (NHS, 37°C), short-term HS (SHS, 42°C, 30min) and long-term HS (LHS, 42 °, 6h) conditions. Interestingly, we found that the chromatin conformation of K562 cells was notably stable in response to short-term HS, while showed some characteristic changes after long-term heat treatment, while chromatin accessibility remained stable during the whole process. At the cell cycle level, we found that chromatin conformation of G1/S cells remained largely stable throughout the whole process of HS, while G2/M cells changed slightly after LHS. Based on these data, a cell cycle-stage redistribution model, i.e., the accumulation of cells at M stage in response to LHS, emerged, and it was further supported by *in silico* and experimental evidence.

Our results provided a plausible cellular mechanism, allowing cells to adapt to long-term thermal stress, and highlighted the distinct mechanism of gene regulation in reaction to immediate crisis and environmental adaptation. Last, our analysis demonstrated that cell cycle consideration is essential in the interpretation of bulk Hi-C data of HS and responses to other stimuli.

## Results

### Chromatin conformation remains largely stable after short-term HS, but undergoes substantial changes after long-term heat treatment

To understand the response of chromosomal conformation to short- and long-term HS, we performed Hi-C experiments [19] on human K562 cells under normal culture temperature (37°, NHS), 42° for 30 minutes (short, SHS) and 6 hours (long, LHS) with at least two biological replicates for all conditions (Fig. S1A). The Hi-C library was highly reproducible, given that the median GenomeDisco [30] scores between replicates were at least 0.85 among all chromosomes for all three conditions with bin size larger than, or equal to, 50kb (Fig. S1B). We merged the replicates, which yielded about 490 million (M), 315M and 567M pairs of contacts for NHS, SHS and LHS, respectively, for all subsequent analysis (Table S1).

Hi-C contact maps were highly similar between NHS and SHS, while substantially changed after LSH (Fig. 1A and S1C). First, the overall contact matrices and contact frequency decay curves (also known as p(s) curve [31]) were remarkably similar between NHS and SHS, while very different from those of LHS. The GenomeDisco score between the Hi-C map of NHS and SHS was comparable to the score between the biological replicates (median = 0.91, binsize = 50Kb), while this same score was significantly smaller between SHS and LHS (median = 0.73, binsize = 50Kb, Fig. S1D). Using Jensen-Shannon divergence (JSD) to measure the closeness of two p(s) curves, we found the JSD to be 0.012 and 0.062 between NHS and SHS and between SHS and LHS, respectively (Fig. 1B). The much smaller JSD indicates a relatively stable p(s) curve in response to SHS, while dramatic changes happened after LHS, e.g., increased distal contact frequency from about 0.5Mb to 13.2Mb for LHS (Fig. 1B).

**Fig. 1.**
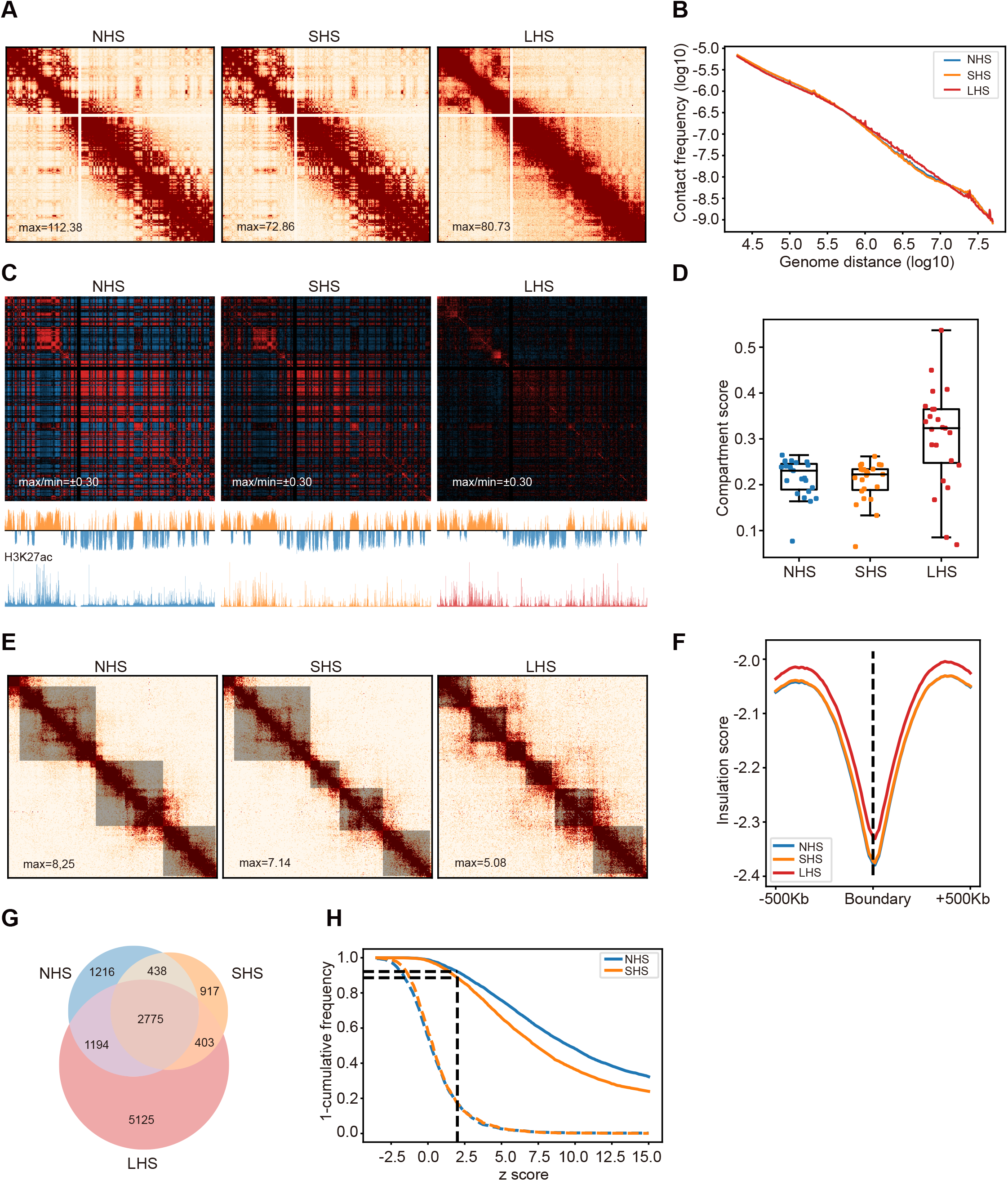
Chromatin conformation remains stable after short-term HS, while it is mostly altered after long-term HS. (A) Chr6 was shown with 500Kb resolution as an example of the contact maps in K562 cells during the process of HS. (B) Contact frequency decay curves at NHS, SHS and LHS conditions. (C) Chromatin compartmentalization at the three conditions. Chr6 was shown as an example. The autocorrelation matrices together with the first eigenvector profiles were shown. In the first eigenvector, compartment B was colored as blue and A as orange. (D) Genome-wide compartment scores at each condition. (E) TADs detected in a 4Mb region centered at the TSS of HSPA1A gene were shown as an example, together with the corresponding contact maps. The detected TADs were shaded. (F) Genome-wide insulation score profiles around TAD boundaries at the three conditions. (G) Numbers and proportions of overlap of chromatin loops at the three conditions. (H) Distributions of z-scores of LHS-specific loops in NHS and SHS conditions. Dashed lines indicated the distributions of z-scores of the randomly chosen bin pairs with the same genome distances. Black dashed line showed the proportion of LHS-specific loops that had z-scores of contacts > 2 compared to their flanking regions.

Second, HS lasting 30 min slightly shrank the A/B compartments [19], while substantially changed the plate pattern in the autocorrelation matrices after LHS (Fig. 1C). Genome-wide, 3.6% of bins underwent compartment switches between NHS and SHS, while more than six-fold more bins (23.04%) switched compartments after LHS (Fig. S1E). Quantitatively, the differences in compartment strengths (AB/AA+BB) between NHS and SHS were minor, as the chromosome-wise average compartment strengths were 0.2143 and 0.2069 (*p* = 0.0014, *t*-test), respectively, while grew to 0.3045 in LHS (*p* = 1.6 × 10 ^−4^, and 5.71 × 10 ^−5^, comparing LHS with NHS and SHS, respectively, *t*-test) (Fig. 1D).

Third, TADsremained largely unshifted in response to HS [32], irrespective of the duration of heat treatment, while became much weaker after LHS (Fig. 1E). Using deDoc [33], we called 2104, 2207 and 2307 TADs in NHS, SHS and LHS at 10Kb resolution, respectively. Comparing the TAD boundaries between any two conditions, at least 40% of boundaries remained stable, not shifting more than one bin (Fig. S2A). Even the altered boundaries did not change substantially, as intra-TAD contacts remained enriched when annotating TADs with those called in another condition (Fig. S2B). The percentage of stable TAD boundaries between conditions was three-fold greater than that between cell lines. For example, only about 11.2% of TAD boundaries were found to be stable between GM12878 and K562 compared to public Hi-C data [21]. The similarity between NHS and SHS was comparable to that between replicated libraries, as about 56% and 54% of boundaries were found stable, respectively. Despite the stability of TAD positions and the similarity of TAD strengths between NHS and SHS, those strengths became much weaker after LHS. The insulation scores (IS) of TAD boundaries [34] in SHS only reflected 0.19% of changes compared with NHS, i.e., with mean IS = -2.3722 and -2.3768 (*p* = 0.0028, paired *t*-test), respectively, while reflected 1.91% changes after LHS (IS = -2.3269, *p* = 1.15 × 10 ^−67^, paired *t*-test, Fig. 1F). Moreover, TAD strength scores (TS, defined as the fraction of inter-TAD contacts in total intra-chromosome contacts) [35] in SHS were only 0.99% different from those of NHS, i.e., chromosome-wise mean TS = 0.3838 and 0.3876 (p = 0.036, paired *t*-test), respectively, while increased 18.45% to 0.4546 at LHS (*p* = 1.23 × 10 ^−19^, paired *t*-test, Fig. S2C). Taken together, we found that TADs could remain stable within 30 mins of HS exposure and largely remain unshifted, but became much weaker after six hours of HS.

Fourth, chromatin loops were less stable in response to the HS; however, the strength and significance of these changes were minor. Using HICCUPS [36], we identified 5623, 4533 and 9497 chromatin loops longer than 200kb initiating from 8150, 6494 and 13781 anchors in NHS, SHS and LHS, respectively (Fig. 1G). Although only 2779 loops were shared by all three conditions (Fig. 1G), the following evidence supports that the alterations of condition-specific loops were minor. Aggregated peak analysis (APA) showed that the contact frequencies between the anchors of condition A-specific loops were also highly enriched in condition B (Fig. S2D) where A and B could be any one of the three conditions. Since the fraction of shared loops between NHS and SHS is nearly comparable to that of shared loops between biological replicates in the public GM12878 dataset [21] (Fig. 1G and S2E), we now mainly focus on loops specific to LHS. Here, we found that the contacts between LHS-specific loop anchors in the NHS and SHS conditions were also significantly higher than those in the flanking region, i.e., 92.16% and 88.62% of LHS-specific loops also had z-scores of contacts > 2 compared to their flanking regions in NHS and SHS conditions, respectively (see Method, Fig. 1H), while only about 17% of such bin pairs could be seen in random control (Fig. 1H). This observation indicated that contacts had been concentrated between these LHS-specific loop anchors in the NHS and SHS conditions. Furthermore, the binding affinities of CTCF to the anchors of LHS-specific loops were nearly comparable with those of all combined loops in NHS (*p* = 0.05, *t*-test, Fig. S2F). Finally, genes located in the anchors of these LHS-specific loops did not enrich any GO [37] functional terms (with threshold q-value <0.05, Table S2). These data collectively imply that the condition-dependent specificity of these loops may be rather weak or may be resulted from the insensitivity of the state-of-the-art chromatin loop callers [38].

Altogether, our data suggested that chromatin conformation may remain stable under short-term HS, but that it could gain notable changes after long-term heat exposure at all levels.

### Local nucleosomal environment remained stable during the whole process of HS

To characterize the local nucleosomal environment, e.g., the accessibility of chromatin, we performed ATAC-seq with two biological replicates [39], obtaining 55, 64 and 54 million de-duplicated reads for NHS, SHS and LHS cells in total, respectively. The data were highly reproducible (Fig. 2A and S3A) and had characteristic peaks in fragment size distribution representing mono-, di-, and tri-nucleosomes (Fig. S3B), indicating high quality.

**Fig. 2.**
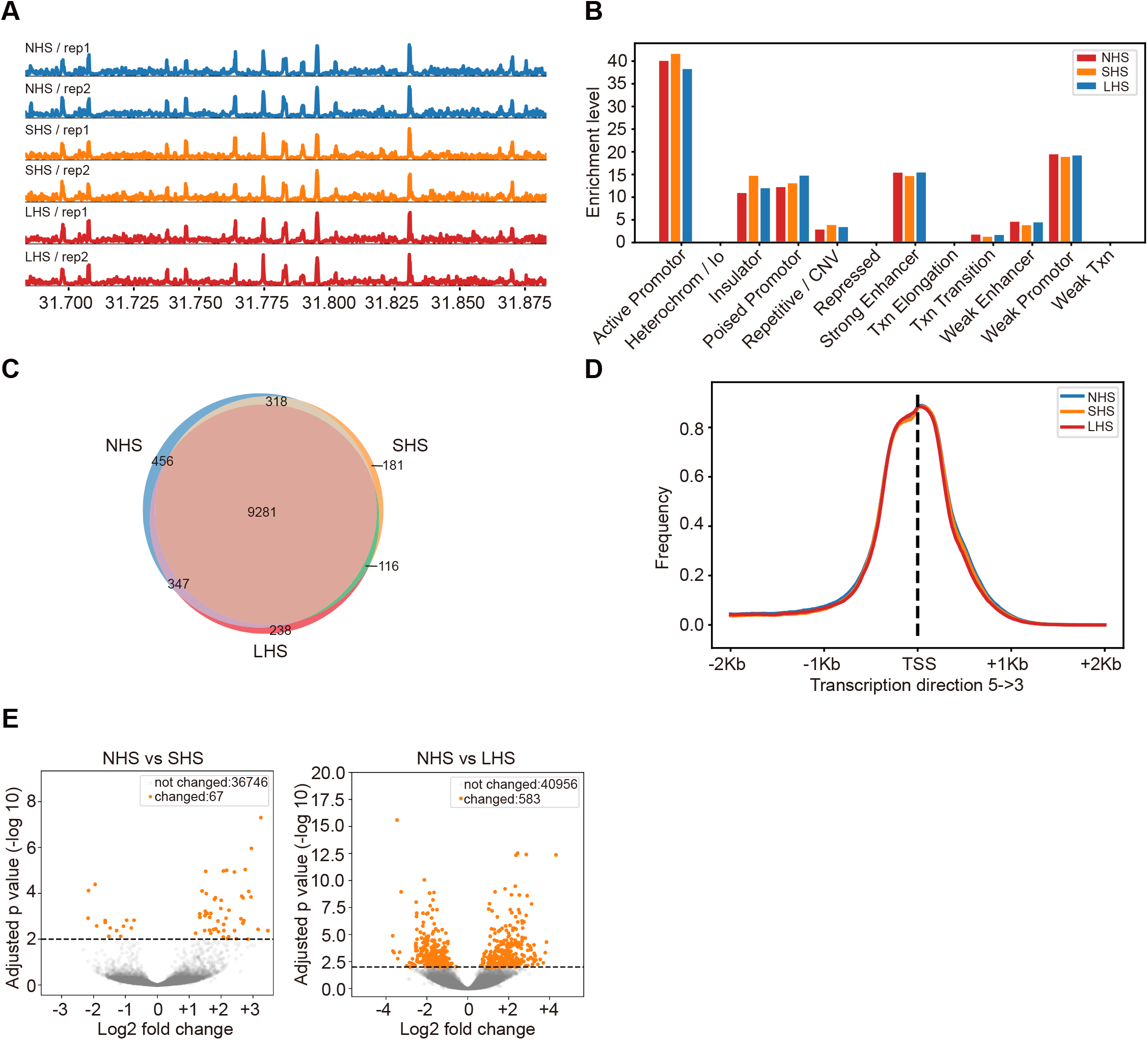
Local chromatin environments remained stable during the whole process of HS. (A) ATAC-seq signal profiles (observed / expected calculated by MACS2) of the biological replicates at the NHS, SHS and LHS conditions. The 200Kb region centered at the TSS of HSPA1A gene was shown. (B) Distribution of ATAC-seq peaks over genome elements in the three conditions. (C) Numbers and overlaps of accessible genes (TSSs covered by NFRs) in the three conditions. (D) Distributions of NFRs around TSSs of accessible genes. (E) Volcano plot showing the results of the differential accessibility analysis in NFRs before and after HS. Accessibility was measured by the coverage depth of short ATAC-seq fragments.

As the indicator for the local nucleosomal environment, the characteristics of ATAC-seq remained stable after HS. First, the ATAC-seq signal profiles were highly correlated with each other in the three conditions with similar cosine measurements larger than 0.95 when comparing any two conditions (Fig. 2A). A total of 30326, 25503, and 26358 recurrent peaks were called for NHS, SHS and LHS by MACS2 [40], respectively. More than 80% of the peaks overlapped with more than one bp between any two conditions, which is a level similar to that between biological replicates (85% to 93%). Notably, 19490 peaks were found overlapped by at least one bp between all three conditions. Less than 5% of peaks were identified as different between conditions by DESeq2 [41] (FC>1 and p<0.01) (Fig. S3C). The distribution of ATAC-seq peaks over genome elements was also found to be nearly identical among the three conditions (Fig. 2B) with most peaks located in the promoters and enhancers. Second, nucleosome-free regions (NFR) in the promoters, as a particularly interesting local chromatin feature, were found to be highly stable in response to heat. Using our nucleosome detection algorithm deNOPA [42], we identified 113429, 90101 and 95576 high-quality NFRs in NHS, SHS and LHS, respectively (Fig. S3D and E). Highly enriched ubiquitin-related GO terms in heat-shock specifically accessible genes (with TSS covered by NFR) further indicated the high quality of the called NFRs [43] (Fig. S3F). Overall, HS had little effect on NFRs. For example, compared to NHS, the percentages of intact NFRs in SHS and LHS (52.50% and 53.20%, respectively; see Method) were comparable to those between biological replicates (55.17% in NHS). Most (84.86%) of the expressed protein-coding genes covered by NFRs in TSS in one condition also contained NFRs in other conditions [44] (Fig. 2C) with almost identical distribution in the TSS regions among conditions (Fig. 2D). The enriched GO terms in these genes were also almost identical among the three conditions (Fig. S3G). Third, the accessibility of chromatin, especially in NFRs, was largely stable during HS. We measured accessibility of the NFRs using coverage depth of short fragments (insertion lengths ≤ 115bp). Results showed that less than 2% of NFRs showed significant accessibility changes after heat stress, even at the loose threshold of FDR<0.05, i.e., 67 and 583 NFRs were changed after SHS and LHS, respectively (Fig. 2E). Collectively, our results showed that the local nucleosomal environment was largely unchanged during the whole process of HS.

### Chromatin conformation of G1/S cells remains largely intact after HS

The cell cycle can be arrested under HS [11, 45, 46]. Consequently, chromatin conformation was significantly different between G1/S and G2/M cells [47, 48]. Therefore, we asked whether alteration of the Hi-C contact map we observed could have resulted from redistribution of cell cycle status in the cell population after LHS. Indeed, we observed a substantially increased percentage of G2/M cells after LHS by FACS analysis (Fig. S4A). Again, we asked how much of the Hi-C map differences we observed could be attributed to this redistribution of cell cycle status, especially since overall local chromatin accessibility was considered stable during the cell cycle [49, 50]. To address these questions, we sorted the cells into G1/S and G2/M phases by FACS in three conditions and performed Hi-C experiments for all sorted cells. The results for G1/S and G2/M cells are presented in this section and the next, separately. For G1/S cells, 236, 616 and 559 million of high-quality contacts were obtained for NHS, SHS and LHS conditions, respectively, with high reproducibility between the biological replicates (Fig. S4B and Table S1).

The overall Hi-C contact maps of G1/S cells are almost identical after SHS and LHS (Fig. 3A). The chromosome-wise average GenomeDisco scores between any two conditions were larger than 0.85, which is much higher than the scores between NHS and LHS in mixed cells (*p* < 1 × 10 ^−10^, paired *t*-test, Fig. S4C). The p(s) curves of G1/S cells were also similar among the different conditions (Fig. 3B), considering that the JSD scores of two p(s) curves between two temporally sequential conditions were 0.038 and 0.023, respectively, both much smaller than the JSD between SHS and LHS in mixed cells. From the p(s) curve, little increase of ultra-long interactions (>10M) was noted after HS, which is also revealed by the fold change of contact profiles (Fig. S4D), but the change is slight compared with that of mixed cells (Fig. S1C).

**Fig. 3.**
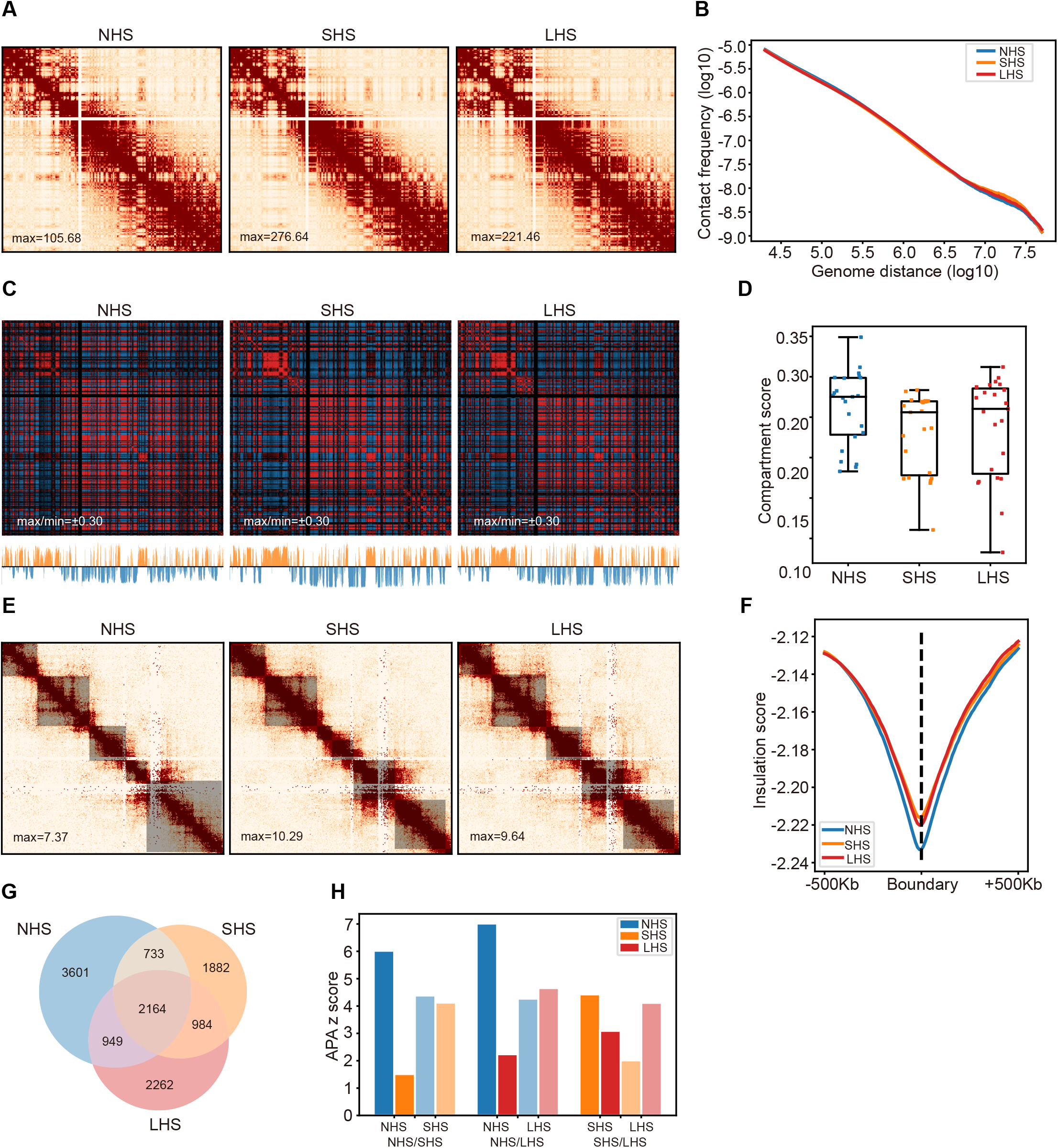
Chromatin conformation in G1/S cells remains largely intact after long-term HS. (A) An example showing the contact maps of G1/S phase cells in the process of HS; chr6 was shown with 500Kb resolution as in Figure 1A. (B) The contact frequency decay curves at the three conditions in G1/S phase cells. (C) Chromatin compartmentalization at the three conditions in G1/S phase cells. Chr6 was shown as an example as in Figure1C. The autocorrelation matrices, together with the first eigenvector profiles, were shown. (D) The chromosome-wise compartment scores at each condition in the G1/S phase cells. (E) TADs detected in the same region as that shown in Figure1E with G1/S cells, together with the corresponding contact maps, were also shown. (F) Genome-wide insulation score profiles of G1/S cells around TAD boundaries at the three conditions. (G) Numbers and proportions of overlap of chromatin loops at the three conditions for G1/S cells. (H) Comparison of APA score (observed / expected KR normalized contact frequencies) among the different conditions in G1/S phase cells. For each comparison, the APA results of loops specific to first (darker color) and second (lighter color) conditions were calculated using the contact frequencies in the corresponding two conditions marked by different colors.

The A/B compartment patterns were much more stable in G1/S cells than those in mixed cells. None of alterations observed in mixed cells, e.g., shrinkage of the autocorrelation matrix and the large proportion of compartment switching, was observed in G1/S cells (Fig. 3C and S4E). Compared to the near 0.09 increase of average compartment strength in the mixed cells after long-term HS (Figure 1D), the score changes in any comparisons with G1/S cells were smaller than 0.04 (p < 10 ^−5^, paired *t*-test, Fig. 3D). A much smaller percentage of genome regions had also switched compartments after SHS and LHS, i.e., 11.3% and 6.1% in G1/S versus 3.6% and 23.04% in mixed cells, respectively (Fig. S1E and S4E).

G1/S cells showed intact TAD structure during HS (Fig. 3E). We identified 2684, 2846 and 2713 TADs in NHS, SHS and LHS with median lengths of 740Kb, 750Kb and 780Kb, respectively, in G1/S cells. Compared to about 40% in mixed cells (Figure S2A), about 50% of TAD boundaries were shared between any two conditions in G1/S (Fig. S5A and B). Interestingly, both IS and TAD scores were slightly elevated after SHS (*p* = 4.9 × 10 ^−6^ and 4.05 × 10 ^−11^, respectively, paired *t*-test) and remained stable for IS (*p*= 0.20, Fig, 3F), or slightly decreased for the TAD score (*p*= 5.43 × 10 ^−7^, Fig. S5C), during LHS. However, the magnitude of the changes in both IS and TAD scores was significantly smaller than that between NHS and LHS in the mixed cells (Fig. 1E and F).

Loop structures were also largely stable after HS in G1/S cells with fewer condition-specific loops found compared to mixed cells. There were 7445, 5761 and 6357 intra-chromosome loops called in NHS, SHS and LHS conditions separated by more than 200Kb, respectively (Fig. 3G). Among these, 2164 loops were shared among different conditions, and the proportion of common loops between any two conditions was comparable to that between biological replicates in the public GM12878 dataset (Fig. 3G and S2E). In addition, for each condition, the contact frequencies were also high between loop anchors called in other conditions, indicatin little condition specificity of loops (Fig. 3H and S5D).

Altogether, the overall consensus of p(s), A/B compartments, TADs and loops showed a rather stable pattern between SHS and LHS, and slight, or no, changes compared to NHS for G1/S cells.

### Chromatin conformation of G2/M cells changed in the same direction as mixed cells, but in smaller degree

For sorted G2/M cells, two biological replicates were conducted with high reproducibility, and 477, 488 and 623 million valid contacts were obtained from NHS, SHS and LHS, respectively (Fig. 4A and S6A). The overall Hi-C contact maps of G2/M cells experienced similar, but fewer, changes after SHS and LHS compared to those of mixed cells (Fig. 4A and S6B).

**Fig. 4.**
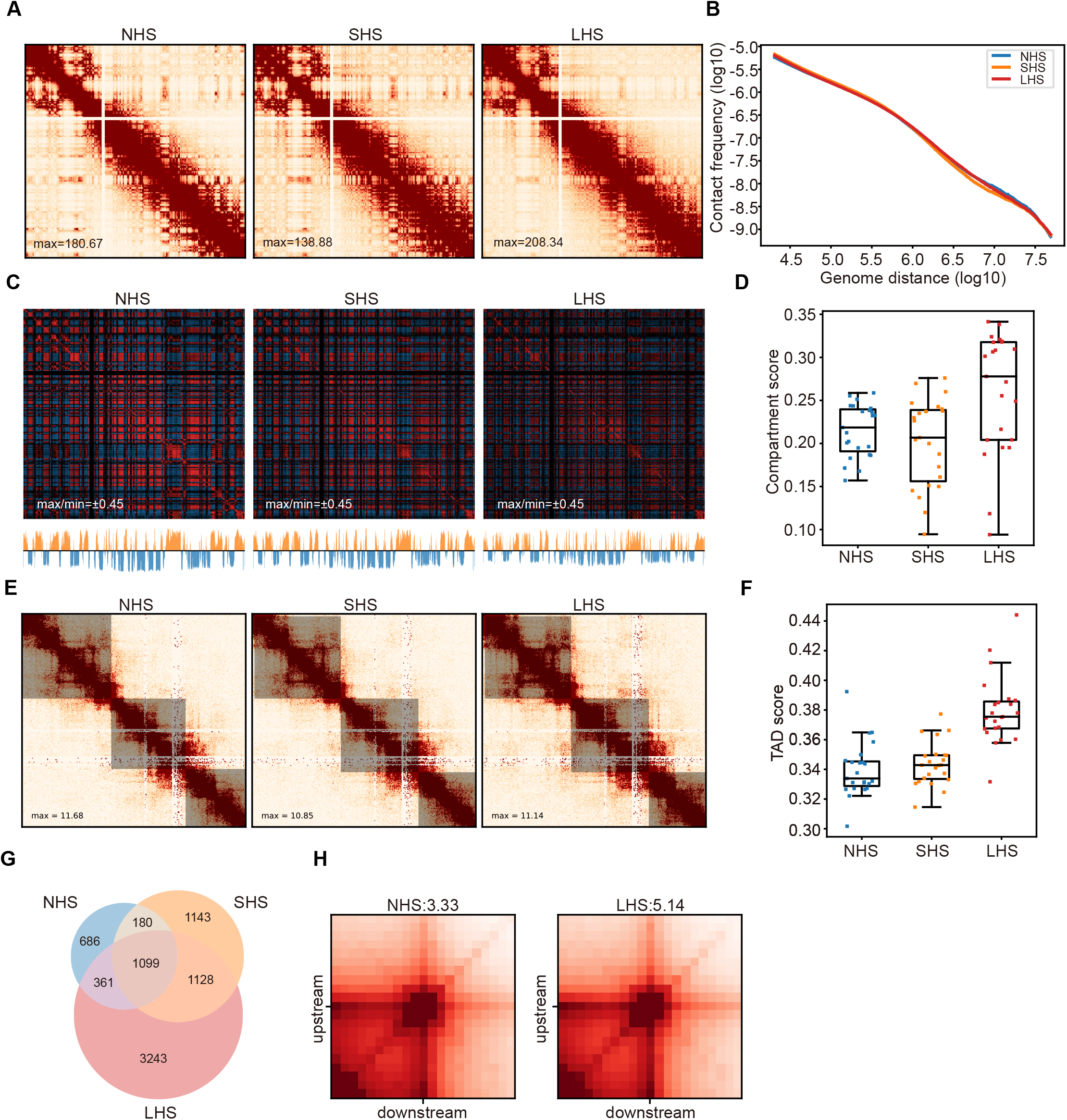
Chromatin conformation of G2/M cells changed slightly during long-term HS. (A) Contact maps in G2/M phase cells before and after HS. Chr6 was shown at 500Kb resolution as in Figure1A. (B) Contact frequency decay curves in G2/M phase cells. (C) Chromatin compartmentalization in G2/M phase cells. Chr6 was shown as an example as in Figure1C. Autocorrelation matrices, together with the first eigenvector profiles, were shown. (D) Chromosome-wise compartment scores in G2/M phase cells in each condition. (E) TADs detected in same region as that shown in Figure1E were shown as an example for G2/M cells, together with the corresponding contact maps. Detected TADs were shaded. (F) TAD strengths indicated by TAD scores in G2/M phase cells at the three conditions. (G) Numbers and proportions of overlap of chromatin loops in G2/M phase cells. (H) APA scores for the LHS-specific loops are shown in NHS and LHS conditions.

The Hi-C contact matrices of G2/M cells were similar between NHS and SHS and changed somewhat in LHS (Fig. 4A), i.e., median GenomeDisco scores were 0.83 and 0.81 between NHS and SHS and SHS and LHS, respectively (Fig. S6B), which is larger than that of scores in the mixed cells after long-term HS (0.74 and 0.72, when comparing LHS with NHS or SHS, respectively). Similarly, JSD score of p(s) curves between SHS and LHS in G2/M cells was 0.040, smaller than that in mixed cells (0.062) (Fig. 1B, Fig. 4B and Fig. S6C).

The A/B compartment of G2/M cells changed during LHS, while the changes were smaller than those of mixed cells. The plate pattern of autocorrelation matrix also shrank at LHS for G2/M cells, but to a lesser degree than that in mixed cells (Fig. 4C and 1C). Quantitatively, compartment scores were comparable between NHS and SHS (with chromosome-wise average compartment scores 0.22 and 0.20, *p* = 0.03, paired *t*-test), but elevated significantly in LHS (0.26, *p* = 3.14 × 10 ^−9^, Fig. 4D). However, after LHS, the compartment score was still smaller than that of mixed cells, indicating stronger compartmentalization than that in mixed cells (0.30, *p* = 0.02 paired *t*-test). In addition, fewer chromatin regions switched compartments between any two conditions in G2/M cells compared with mixed cells (< 8.5%, Fig. S6D).

TAD structure of G2/M cells also remained intact in SHS, but slightly changed in LHS, albeit to a minor degree in comparison to that in mixed cells (Fig. 4E, S7A and S7B). Quantitatively, the chromosome-wise average TAD scores were 0.3397 and 0.3437 in NHS and SHS, respectively (*p*= 0.10, paired *t*-test), while increased to 0.3796 after LHS (*p*= 1.16 × 10 ^−12^, Fig. 4F), which was still smaller than that found in mixed cells (0.4546, *p* = 2.08 × 10 ^−16^).

Similar to mixed cells, some weak alterations were found at loop level. There were 2326, 3550 and 5831 loops called in NHS, SHS and LHS for G2/M cells with distances greater than 200Kb between anchors, respectively. 1099 loops were found overlapped among the three conditions (Fig. 4G). Similar to the mixed data, the APA scores for the condition-specific loops were only slightly increased compared to the scores calculated in the condition not regarded as loops (Fig. 4H and S7C). Further, few differences of CTCF-binding strength (Fig. S7D), or heat-shock-related GO terms enrichment (Table S3), were found for LHS-specific loop anchors.

Taken together, the trends of chromatin conformation changes for mixed and G2/M cells were all similar although the degree of such changes was smaller for G2/M cells.

### Redistribution of G1/S and G2/M cells in response to HS explains Hi-C contact map changes after LHS

Given the rather stable chromatin conformation observed in the G1/S and G2/M cells in response to HS, it was natural to speculate that the changes observed in the mixed cells stemmed from the changes of cell composition of cell cycle *per se*. By FACS, we found that the percentage of G2/M cells changed from 6.70% in NHS to 5.54% and 17.62% in SHS and LHS, respectively (Fig. S4A). Here, we asked whether cell cycle arrest might explain the observed Hi-C contact map changes.

First, the alteration of chromatin conformation from NHS to LHS mimicked the transition from G1/S to G2/M. Conformation was reported to be different between interphase and metaphase cells [48], in which the M phase cells were characterized by a slow decrease in p(s) curve between 100KB and 10 Mb, together with loss of compartments and TAD structure [47]. Similarly, in our data, we observed the same pattern when comparing LHS to SHS with NHS according to p(s) curves (Fig. 1B). We also observed weaker compartmentalization and TAD strength in LHS (Fig. 1C, 1D, 1F and S2C), and the increased proportion of the M phase cells in G2/M cells coincided with chromatin conformation changes of G2/M cells during the heat-shock process. To quantify the proportion of M cells in G2/M, we manually counted M cells from FACS-sorted G2/M cells by microscope (see Method). In total, we manually counted 343, 285 and 409 G2/M cells in three independent inspections and identified an average of 34.7%, 36.8% and 47.7% M cells in NHS, SHS and LHS, respectively (Fig. 5A and Table S4). Further, the Hi-C contact maps of mixed cells seemed to be an average of those of G1/S and G2/M cells. The p(s) curve of mixed cells was found to be located between the curves of G2/M and G1/S cells, and TAD length also showed the same distributions (Fig. S6C and S8).

**Fig. 5.**
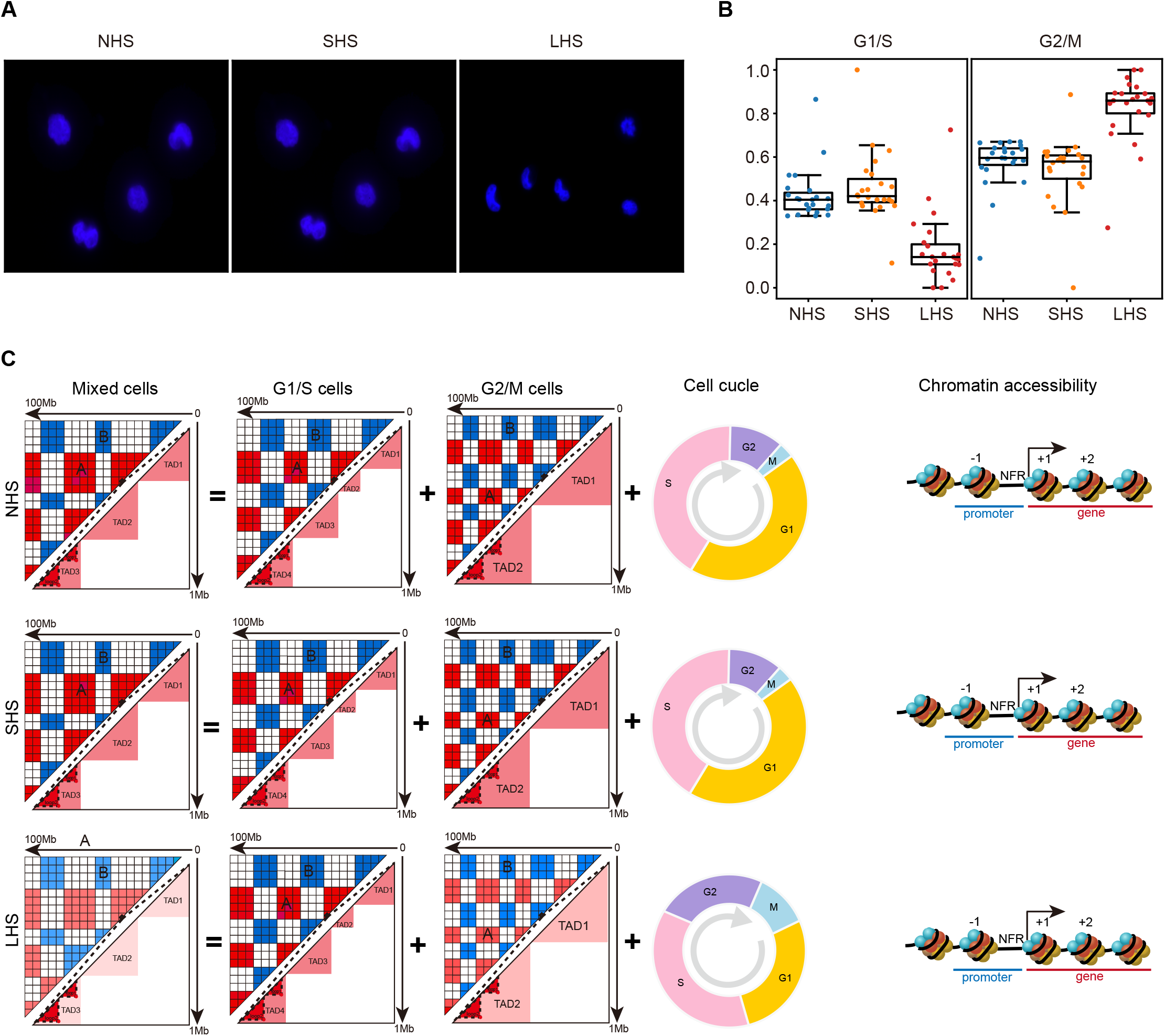
Cell cycle redistribution after LHS contributes to the observed Hi-C contact map changes. (A) Representative images showing cell cycle phase in the sorted G2/M cells for the three conditions. (B) Deconvolved normalized cell fractions of G1/S and G2/M phase cells in the bulks before and after HS and tested by the proportions calculated from different chromosomes. (C) Schematic diagram shows the main discoveries of our study.

Second, we conducted a simple linear deconvolution method to deduce the proportion of cell cycle states from the mixed Hi-C map. Briefly, the Hi-C contact maps were first transformed into distance matrices [51] and then a linear model with matrices in G1/S and G2/M cells as independent variables and those of mixed cells as response variables (Method). The transformation was designed to make the distance matrices independent of sequencing depth to avoid calculating the number of reads per cell. By applying this method to the three conditions, we found that the proportion of G1/S cells remained largely intact in SHS (p=0.15, paired *t*-test, Fig. 5B). After long-term HS, though, the proportion of G1/S cells was found to be dramatically decreased as they deconvolved from most chromosomes (*p* < 10 ^−8^, paired *t*-test). In contrast, the proportion of G2/M cells was found to have dramatically increased as they deconvolved from most chromosomes (Fig. 5B). This trend of cell fraction changes during the process of HS calculated from the Hi-C data was consistent with our FACS analysis. (Fig. S4A).

This evidence strongly points to a model in which redistribution of cell cycle plays a vital role in the observed changes of chromatin conformation in response to long-term HS in mixed cells (Fig. 5C).

## Discussion

In this study, we explored how cells responded to short- and long-term HS with changes in chromatin organization and accessibility. In agreement with the report of Ray and colleagues on K562 cells [26], we observed stable chromatin organization in short-term HS (SHS), which was inconsistent with the reported chromatin remodeling in fly and yeast [23, 24]. This discrepancy may not be completely explained by species-specific response as notable chromatin loop remodeling in response to SHS was also reported in human embryonic stem cells [25]. To large and homothermal animals like human, the cells may be much more heterogeneous in tolerance to temperature changes than the cells of fly and yeast, as heat-shock sensitivity is cell type-dependent [11, 52]. Thus, further studies may be needed to comprehensively survey mammalian cell types to profile the landscape of cellular response to HS in chromatin structure remodeling.

For long-term HS, we observed substantial changes in chromatin structure for mixed cells. The conformation of G1/S cells largely remained stable, and G2/M cells only slightly changed in structure during the whole process of HS. We proposed and showed evidence to support a model whereby conformation changes of mixed cells occurred through the redistribution of cells in cell cycle stages (Figure 5B). We did not provide evidence supporting stable chromatin conformation at each individual cell stage, such as G1, S, G2, and M. Therefore, although we could not completely exclude the possible existence of significant changes at individual cell stages, it is rather unlikely, considering the principle of parsimony. G1/S mixed cells were found to have stable Hi-C maps during HS (Figure 3), and the difference between G1 and S was reported to be minor [48]. If G1 and/or S cells experienced significant changes in chromatin conformation, a rather complicated regulation would need to be applied in order to achieve this overall stability. While only a minor change was observed in G2/M mixed cells (Figure 4), our simple simulation with the equally simple assumption of stable G2 and M conformation showed that such change could be explained by the accumulation of M cells, as also directly observed through imaging. Finally, unchanged chromatin accessibility during the whole heat process for mixed cells further supported our model (Figure 2). Genome-wide assessment of chromatin accessibility using DNase-seq, or ATAC-seq, has shown that mitotic chromosomes preserve significant levels of chromatin accessibility [49, 50]. Thus, if further study were to reveal that chromatin structure at either cell stage changes significantly in response to long-term HS in K562 cells, it would be a result worth investigating in the future.

Stable chromatin structure raises an interesting question in relation to the mechanism of gene repression in response to HS. At the molecular level, it was reported that Pol II pause-release is the step at which the gene’s response to stress is defined [3]. The release of paused Pol II into elongation is inhibited and accumulates at the promoter-proximal pause site of heat-repressed genes in mammals [16]. In addition, changes in expression are accompanied by changes in chromatin environment, including nucleosome dynamics [53], chromatin sumoylation [17], and histone acetylation [16]. Accordingly, it would be interesting to see the overall epigenetic dynamics in the HS process, particularly SHS. SHS might more readily respond to environmental stimulation with chromatin loops, which typically react in minutes owing to fast loop extrusion [54], as compared to global alteration of epigenetic changes, which may react more slowly [55]. It also remains unclear if the accumulation of arrested cells in M stage after LHS is an adaption to heat environment or an indication of the middle stages of transition to apoptosis. At least in our FACS inspection, apoptosis was inconsequential (Fig. S4). Prolonged heat treatment may be needed to answer this question, and it is also subject to the caution of cell type specificity. Collectively, then, our results reached some consensus on the cellular response to HS and also highlighted the necessity of considering cell cycle in interpreting observed changes of chromatin conformation with mixed Hi-C.

## Methods

### Cell culture and HS treatment

K562 cells were maintained at 37°C under 5% CO2 in RPMI-1640 medium (Sigma) supplemented with 10% FBS, 2mM L-glutamate, and streptomycin/penicillin. For short-term HS, growing cells were placed in a water bath for 30 min at 42 °C. For long-term HS, the cell incubator was adjusted to 42° one night in advance, and then cells were placed in a cell incubator at 42° for 6 hours. Cells were brought to 25°C by mixing with one volume of 4°C RPMI-1640 medium and crosslinking immediately.

### In situ Hi-C library preparation

In situ Hi-C was conducted according to the literature [56]. Briefly, K562 cells were cross-linked in 1% formaldehyde solution for 10 minutes and then quenched with 0.125 M glycine. Then, cells were lysed and digested with the MboI restriction enzyme (NEB, R0147). Biotin-14-dATP was used to mark the DNA ends, followed by proximity ligation in intact nuclei. After crosslink reversal, samples were sheared to a length of ∼300 bp and then treated with the End Repair/dA-Tailing Module (NEB, E7442L) and Ligation Module (NEB, E7445L), following the manufacturer’s instructions. Then, biotin-labeled fragments were pulled down using Dynabeads MyOne Streptavidin C1 beads (Life Technologies, 65602). Finally, the Hi-C library was amplified for about 10 cycles of PCR with the Q5 master mix (NEB, M0492L), following the manufacturer’s instructions. Size selection was performed with AMPure XP beads, quantified and sequenced on an Illumina HiSeq X Ten instrument with 2 ×150bp reads.

### ATAC-seq library preparation

ATAC-seq was performed as previously described with a few modifications [57]. Briefly, approximately 50,000 fresh cells were resuspended in 50 μl of ATAC-seq resuspension buffer (RSB; 10 mM Tris-HCl, pH 7.4, 10 mM NaCl, and 3 mM MgCl2) containing 0.1% NP40, 0.1% Tween-20, and 0.01% digitonin and incubated on ice for 3 min. After lysis, 1 ml of ATAC-seq RSB containing 0.1% Tween-20 (without NP40 or digitonin) was used to wash the nuclei. The nuclei were washed and resuspended in 50μl of transposition mix (10μl 5XTTBL, 5μl TTE Mix V50, and 35μl water) and mixed by pipetting up and down 20 times (Vazyme TD501). Then, the transposition reactions were incubated at 37 °C for 30 min in a thermomixer. After “tagmentation”, the sample was purified using Ampure XP beads. The ATAC-seq library was amplified for 11 cycles of PCR with TAE mix (Vazyme TD501), following the manual. The amplified DNA was purified with size selection, quantified and sequenced on an Illumina sequencing platform.

### Cell cycle analysis

2M cells were collected and crosslinked by 1% formaldehyde solution. Then, samples were washed, resuspended with 500ul PBS containing 2% FBS, and permeabilized with 0.1% Triton X-100 for 20 minutes. Then, 30ul 1mg/ml propidium iodide (PI) solution and 4ul 10mg/ml RNAse A were added to the samples. After incubation at 37 °C for 30 minutes, cells were analyzed by flow cytometry.

### Immunofluorescence analysis

About 50,000 cells were fixed with 1% formaldehyde and permeabilized with 0.1% Triton X-100. Then, nuclear counterstaining was performed with DAPI for 5 min at RT. The coverslips were then mounted onto glass slides and sealed with nail polish. Images were acquired with an upright fluorescence microscope (DM5000) with a 100x objective.

### Pre-processing and quality control of sequencing reads

The quality of all libraries was assessed using FasqQC. Reads with a mean quality score less than, or equal to, 30 were removed. For the Hi-C libraries, the 5’-most 15bp of both read 1 and 2 were clipped out because of their low complexity. Adapters were removed by Cutadapt [58], and fragments with length less than, or equal to, 30bp were removed.

### ATAC-seq data processing

All ATAC-seq reads were mapped to human reference genome hg19 using bowtie2 in very sensitive mode with option –X 2000 to restrict the largest fragment lengths from the mapping [59]. The uniquely aligned fragments with mapping qualities smaller than 5 from both sides were filtered out by samtools [60], and duplicates were removed by MarkDuplicates in picard. Signal tracks were built using the bdgcmp subcommand of MACS2 with the “FE” (fold enrichment over control) mode without any control provided [61].

Accessibility for any given region was measured by the number of short fragments (with insertion length bp) with alignment overlapping by one or more base pairs. Differential analysis using DESeq2 [41] was performed to measure the significance of the accessibility changes.

MACS2 peaks were called using the callpeak subcommand of the MACS2 package with the following parameter combination: “callpeak -f BAM -n MACS2 -g hs -q 0.01 --nomodel --shift 100 --extsize 200 --keep-dup all”. Peaks were first called in individual replicates. Then reads from different replicates were merged, and another round of peak calling was performed. Repeated peaks were then taken as those called from the merged reads that overlapped with one of those called in each replicate. The NFR regions were directly called from the merged reads in all replicates using the default configuration of the deNOPA program [57]. Annotations of the regions (peaks or NFRs) were performed using the ChIPseeker package [62] against the RefSeq protein coding genes. The promoters were taken as the interval between the 2Kb upstream and 500bp downstream of the TSSs.

The similarity for any pair of region sets (peaks or NFRs, denoted as A and B) was defined as 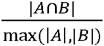, where | * | denoted the number of regions in the set, andA ⋂ B meant regions in the two sets with intersection size larger than 50% of the two.

### Hi-C data processing

Hi-C reads were processed using the HiC-Pro pipeline [58]. Contact reads mapping to ChrY and ChrM or with MAPQ = 0 were filtered out. The p(s) curves were calculated from the genome distances 20Kb to 50Mb separated into 500 bins logarithmically. Contacts were then converted into the ‘.hic’ format used for juicer tools [36]. An A/B compartment profile was called by analyzing the first eigenvector of the KR normalized contact maps at 100Kb resolution [19]. The compartment with higher H3K27ac ChIP-Seq signals was defined as compartment A. TADs were called using deDoc at 10Kb resolution with default configuration [33]. Loops were called using the ‘hiccups’ subcommand of juicer tools with the following parameter combination: ‘-m 2048 -r 10000 -k KR -f 0.01’. Aggregated peak analysis was conducted using the apa subcommand of juicer tools in 10Kb resolution with the following parameter combination: ‘-n 20 -w 10 -r 10000 -k KR -u’. Loop differential analyses were conducted based on the called loops using the hiccupsdiff subcommand of juicer tools with default configuration.

The similarity between any TAD boundary sets B_1_ and B_2_ was defined as the proportion of the boundaries in B_1_ with distance to the nearest one in B_2_ less than 20Kb. Two loops were considered the same if and only if the positions of their anchors were exactly the same.

The z score of a loop was calculated as follows. Suppose the contact matrix was denoted as {c_i,j_}. Then for a loop linking bins i and j, the z score was defined as

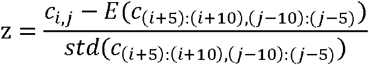

where E and std stood for the mean and standard deviation, respectively. Briefly, the deconvolution was conducted as follows. The contact maps of each chromosome were normalized by matrix balancing [63]. The normalized contact maps were then converted into the distance matrices following the description in [51]. A linear model was built using the distances at G1/S and G2/M stages as independent variables and mixed cells as dependent variable for each chromosome, and the normalized regression coefficients were then reported as the cellular proportions.

## Declarations

### Data and code available

These data were deposited in the Genome Sequence Archive in National Genomics Data Center [64], Beijing Institute of Genomics (BIG), Chinese Academy of Sciences.

### Authors’ contributions

BX, XG and ZZ conceived this project. XG, XL, YJ performed the experiments. BX, XG, FL, and ZZ analyzed data. BX, XG, FL and ZZ prepared the manuscript. All authors read and approved the final manuscript.

### Ethics approval and consent to participate

Not applicable.

### Consent for publication

Not applicable.

## Acknowledgements

We thank Mr. David Martin for English language editorial services.

## Supplementary Figures

**Fig. S1**. Chromatin conformation of K562 cells in the process of HS. (A) Sketch map showing how the experiment was designed. (B) GenomeDisco scores showing the reproducibility of the Hi-C libraries. (C) Fold changes of contact frequencies (observed / expected KR normalized) between any pair of the three conditions. (D) GenomeDisco scores among different conditions, showing the similarity between the two heat-shocked conditions. (E) Proportions of genome regions which switched compartments before and after HS; inner: between NHS and SHS and outer: between NHS and LHS.

**Fig. S2**. Changes of TADs and loops during HS. (A) TAD boundary position changes during the process of HS. The values in row I and column j indicate the proportion of TAD boundaries in I which distanced no more than 1 bin with a boundary in condition j. (B) The stacked contact maps in each condition (row) around TADs determined in other conditions (columns). (C) TAD strength scores (inter-TAD contacts/intra-TAD contacts) of the three conditions. (D) Aggregated peak analysis (APA) results of condition-specific loops. APA results of loops specific to the first condition used contact frequencies in the first and second conditions, as shown in the first two columns. Similarly, results of loops specific to the second condition were shown in the third and fourth columns. (E) The proportion of shared loops between biological replicates in the public GM12878 datasets [21]. (F) CTCF ChIP-seq signals in the anchors of LHS-specific loops and all combined loops in NHS condition. The binding affinities were calculated from the ChIP-seq data in ENCODE [65].

**Fig. S3**. Stability of chromatin states before and after HS, as measured by ATAC-seq. (A) Reproducibility of the ATAC-seq libraries, as measured by the proportion of overlap of MACS2 peaks. Two peaks in different conditions were defined as overlapped if the overlapped region was larger than 50% of both their lengths. (B) Fragment length distributions of ATAC-seq at the three conditions. (C) Volcano plot showing the results of differential accessibility analysis in MACS2 peaks before and after HS. (D and E) Distributions of NFRs in different chromatin states (D) and genetic regions (E). (F) GO terms enriched in genes specifically accessible (with TSSs covered by the NFRs) after short-term (left pattern) and long-term (right pattern) HS. (G) GO terms enriched by genes containing NFR in TSS region at each condition (NHS, SHS and LHS from left to right).

**Fig. S4**. Chromatin conformation in G1/S cells remains largely intact after long-term HS. (A) Cell cycle profiles of K562 cells before and after HS determined by FACS. (B) GenomeDisco scores showing the reproducibility of the Hi-C libraries of G1/S phase cells. (C) GenomeDisco scores showing the similarity among different conditions in G1/S phase cells. (D) Fold changes of contact frequencies (observed / expected KR normalized) between any pair of the three conditions for G1/S cells. (E) Proportions of genome regions which switched compartments before and after HS in G1/S phase cells.

**Fig. S5**. Changes of TADs and loops during HS in G1/S cells. (A) TAD boundary position changes during the process of HS in G1/S phase cells. (B) Stacked contact maps in each condition (row) around TADs determined in other conditions (columns). (C) TAD scores of G1/S cells at the three conditions. (D) Aggregated peak analysis results of loop strengths in G1/S phase cells at each condition (row) using the loops detected in other conditions (columns).

**Fig. S6**. Chromatin conformation changes in G2/M phase cells during K562 HS. (A) GenomeDisco scores between biological replicates indicating the high quality of Hi-C libraries. (B) GenomeDisco scores showing the reproducibility of Hi-C libraries of G2/M phase cells in different conditions. (C) Comparison of contact frequency decay curves among G1/S, G2/M and bulk cells in the three conditions. (D) The proportion of genome regions under compartment switches in G2/M phase cells.

**Fig S7**. Changes of TADs and loops during HS in G2/M cells. (A) TAD boundary position changes in G2/M phase cells during the process of HS. The values in row I and column j indicate the proportion of TAD boundaries in I which distanced no more than 1 bin with a boundary in condition j. (B) The stacked contact maps in each condition (row) around TADs determined in other conditions (columns). (C) Aggregated peak analysis (APA) results of condition-specific loops. APA results of loops specific to the first condition using contact frequencies in the first and second conditions were shown in the first two columns. Similarly, results of loops specific to the second condition were shown in the third and fourth columns. (D) CTCF ChIP-seq signals in the anchors of LHS-specific loops and all combined loops in NHS condition for G2/M cells.

**Fig. S8**. Comparison of TAD lengths in the bulk, G1/S and G2/M data at the three HS conditions.

## Supplementary Tables

**Table S1**. Summary of libraries used in this study.

**Table S2**. GO enrichment results of genes, the promoters of which lie in the anchors of LHS-specific loops in mixed cells. Results of comparison between LHS and both NHS and SHS were listed in different sheets, respectively.

**Table S3**. GO enrichment results of genes, the promoters of which lie in the anchors of LHS-specific loops in G2/M phase cells. Results of comparison between LHS and both NHS and SHS were listed in different sheets, respectively.

**Table S4**. Proportion of M phase cells in G2/M-sorted cells in the three conditions.

