## Supplementary figures and images for "Cell cycle arrest explained the observed bulk 3D genomic alterations in response to long term heat shock for mammalian cells"

### Supplemental figure S1

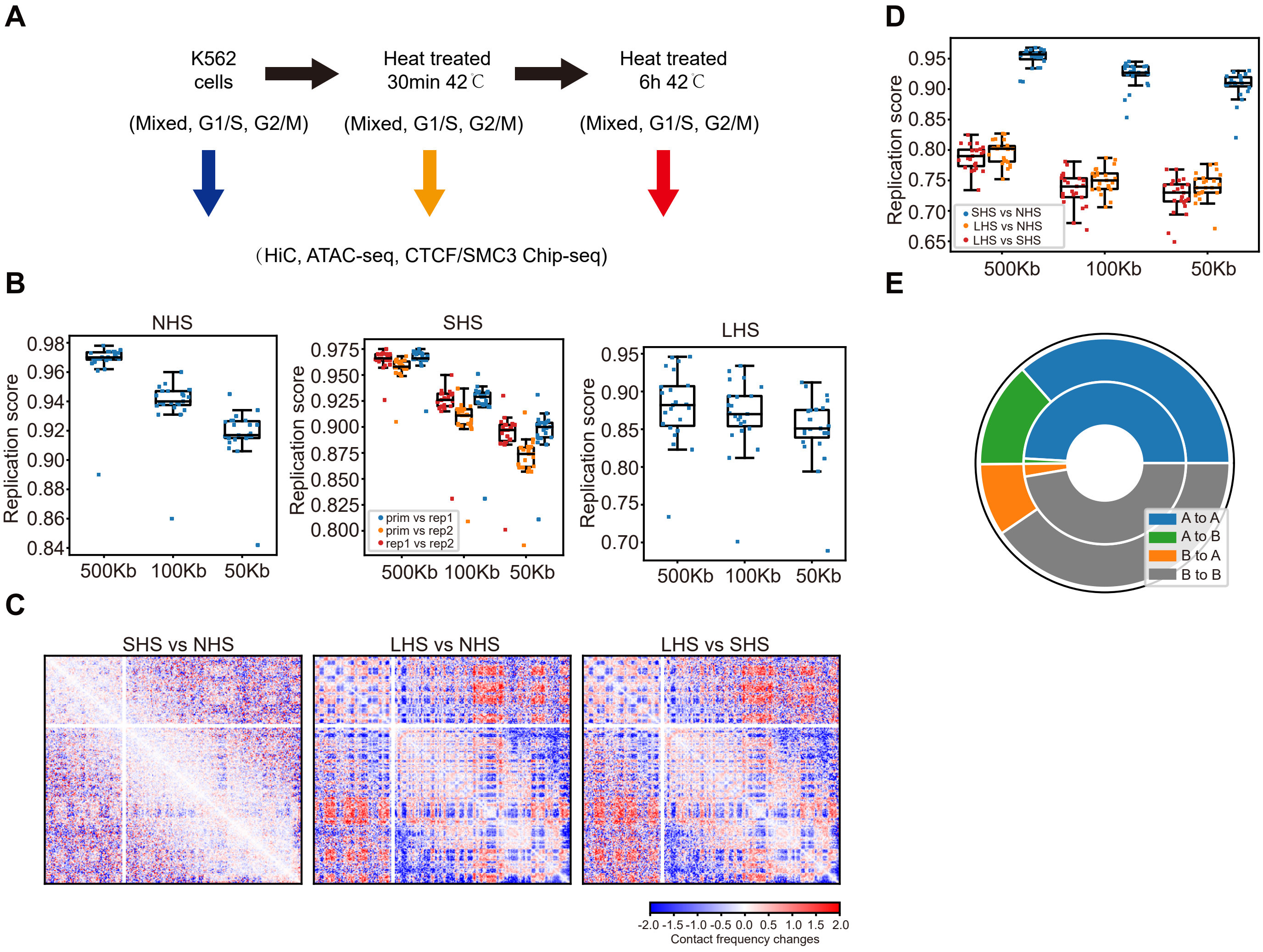

### Supplemental figure S2

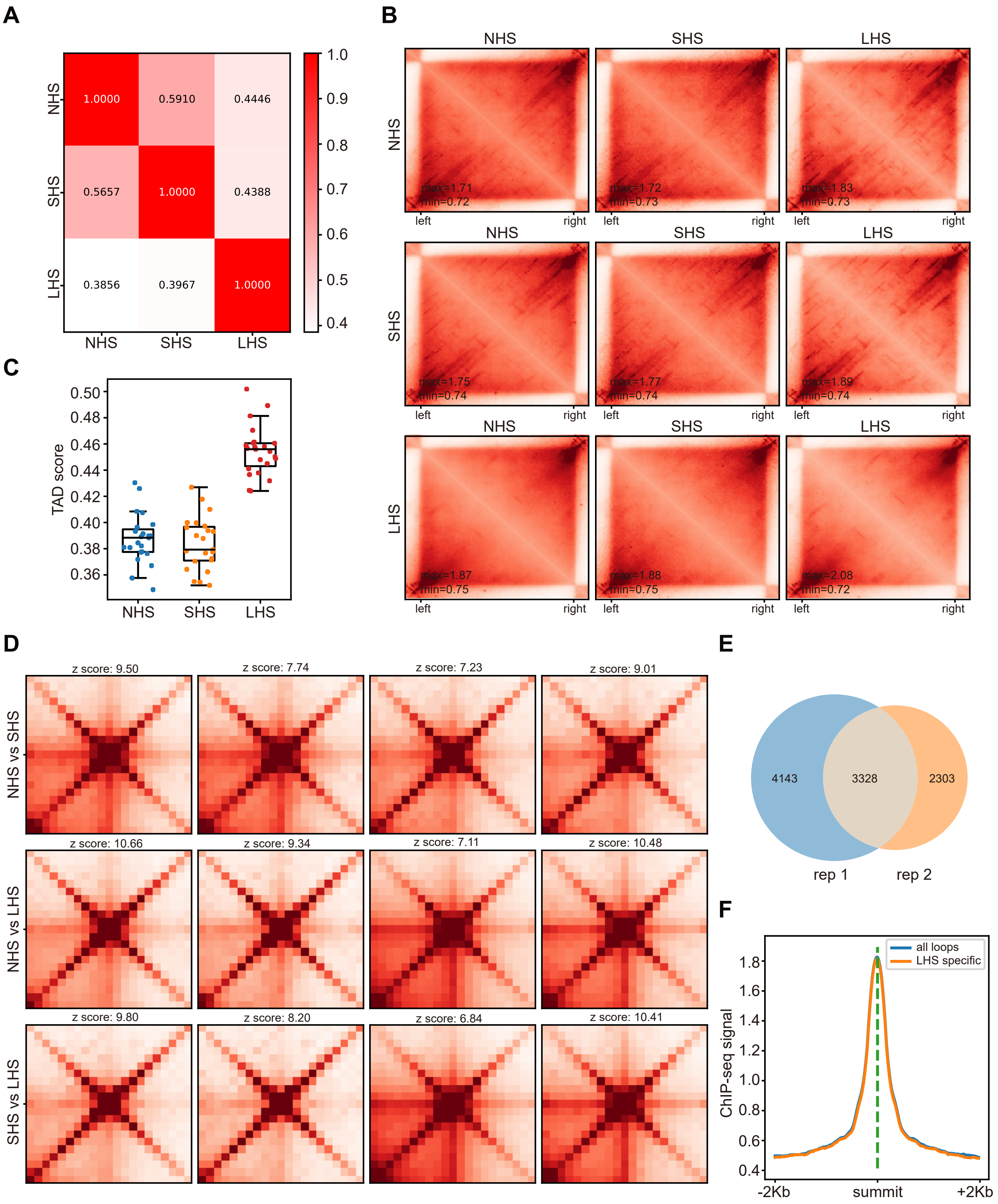

### Supplemental figure S3

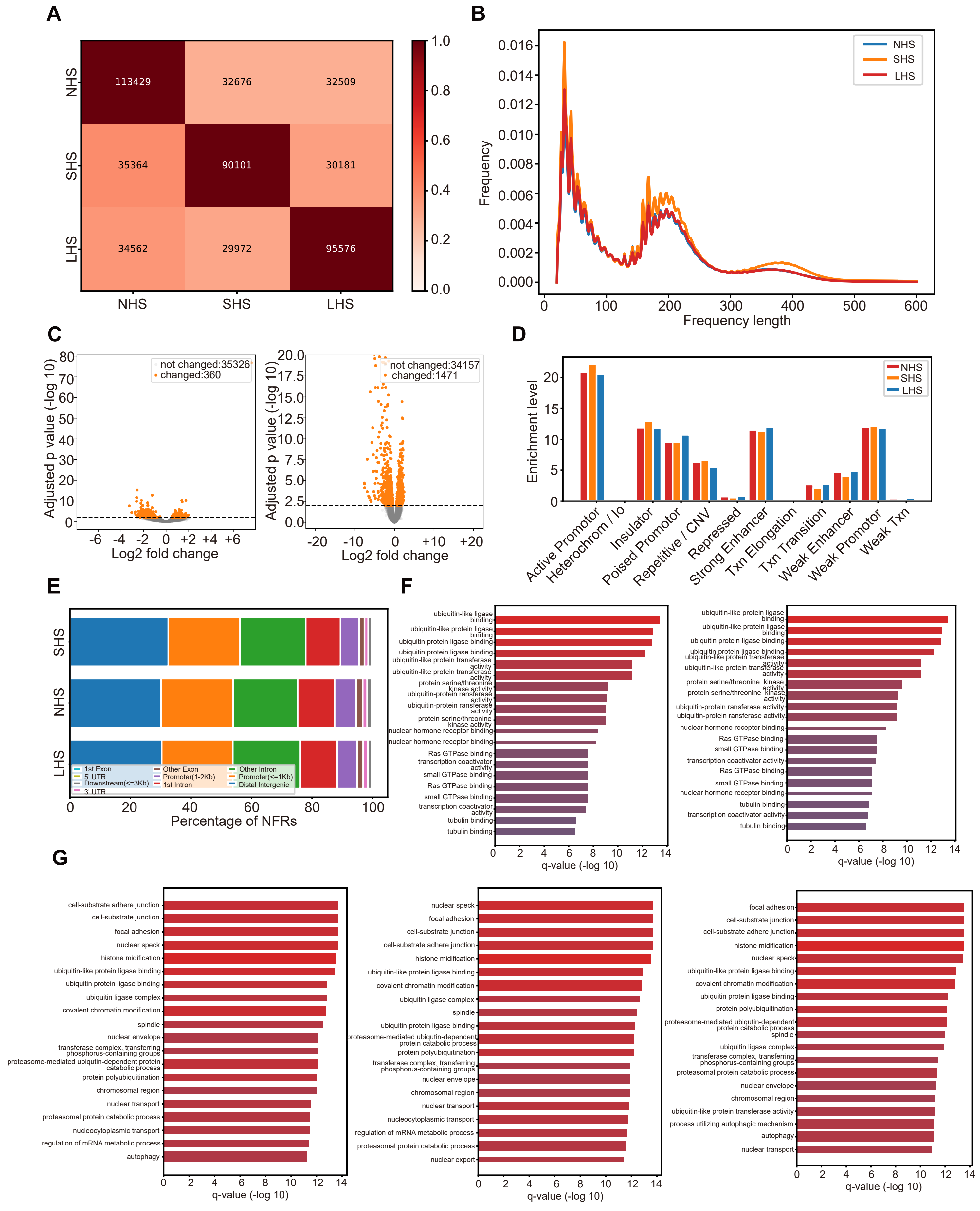

### Supplemental figure S4

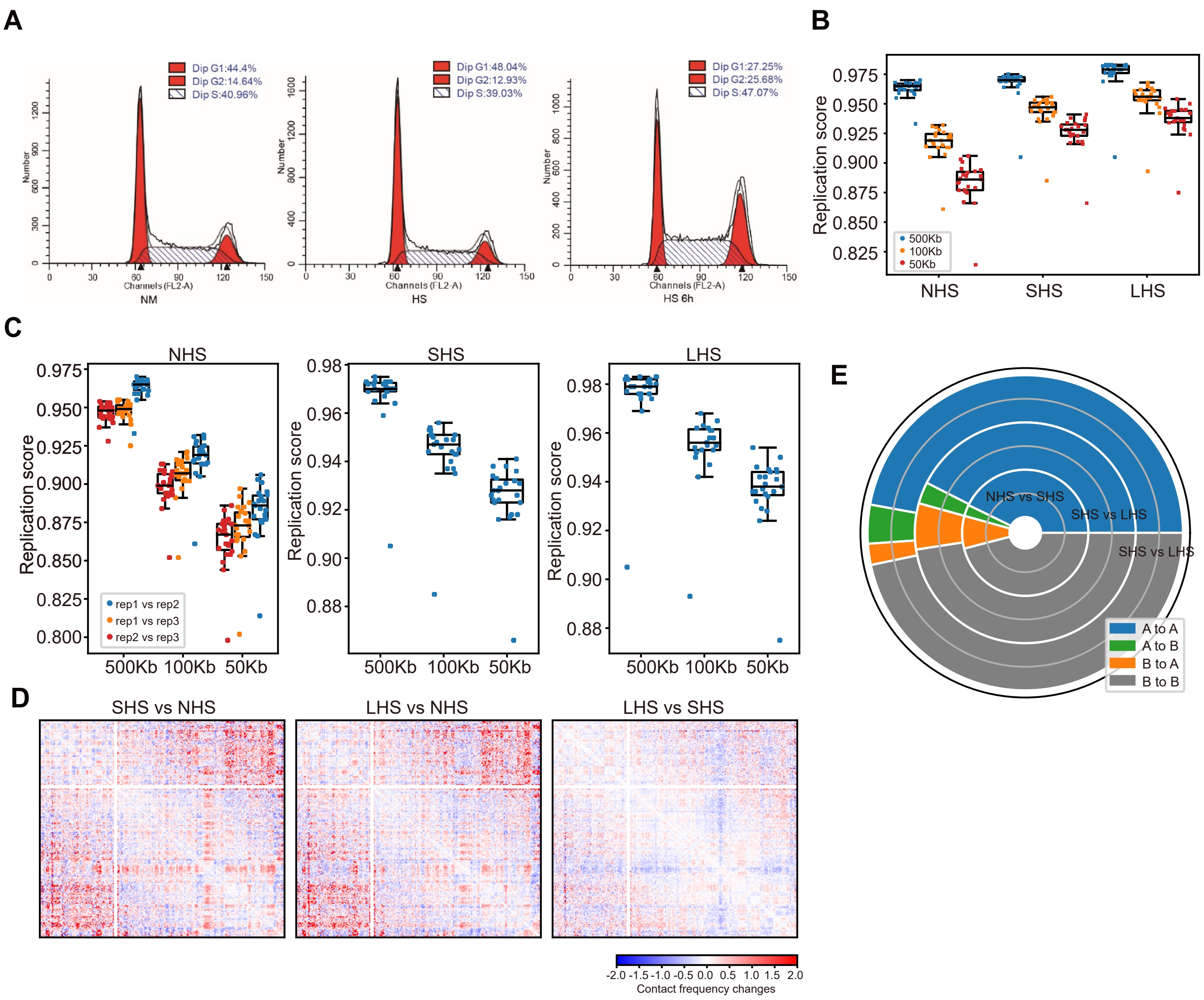

### Supplemental figure S5

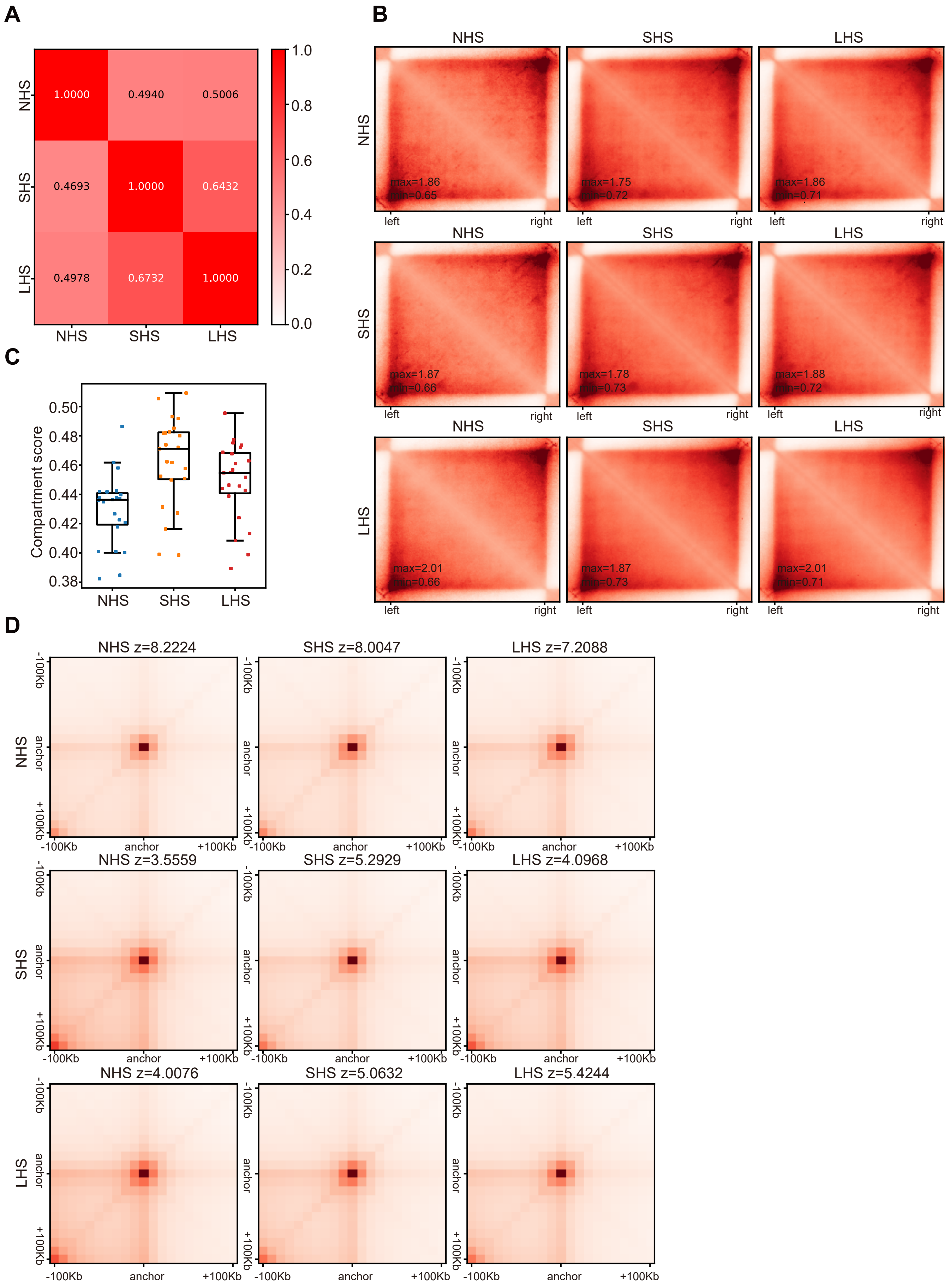

### Supplemental figure S6

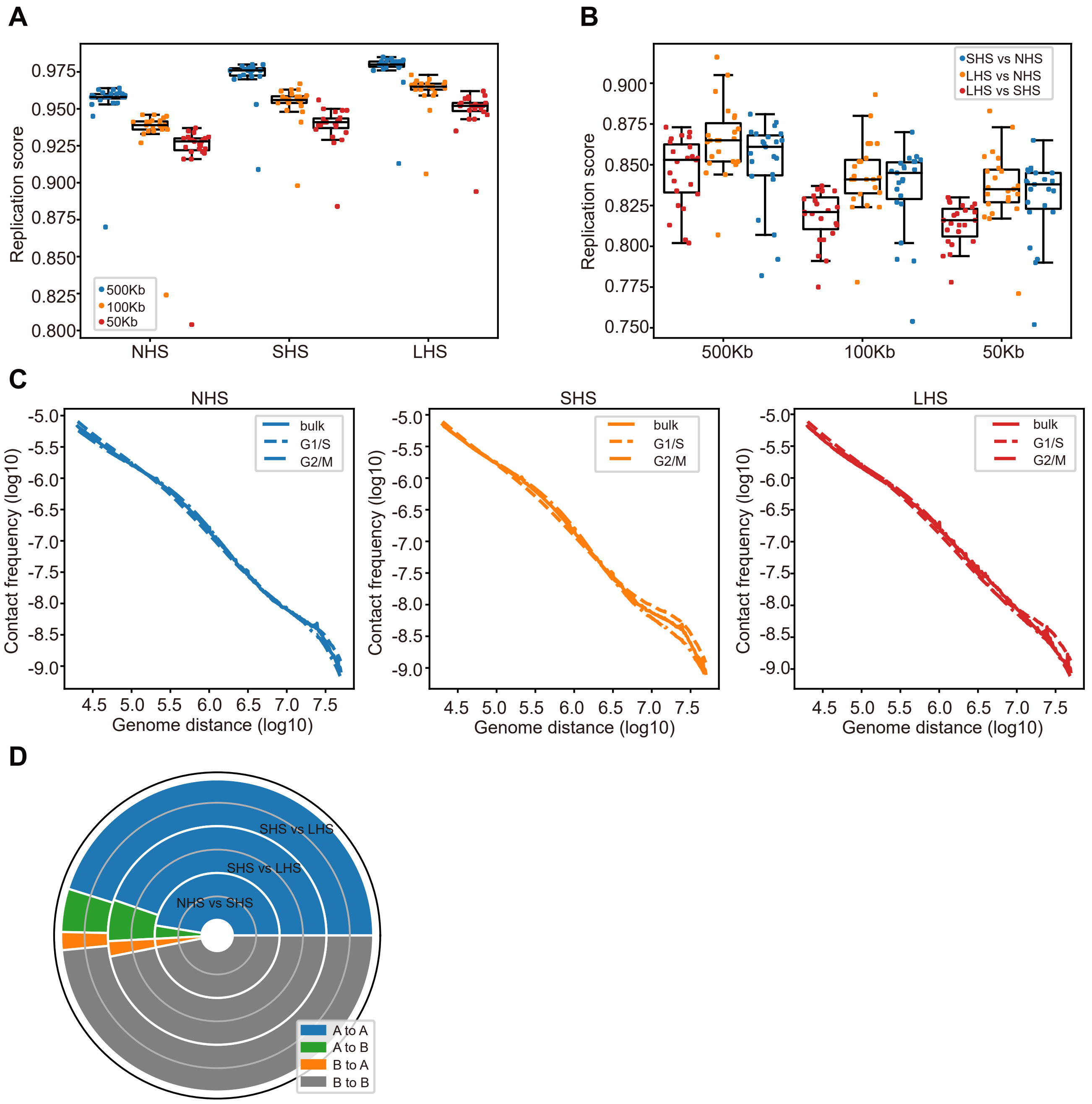

### Supplemental figure S7

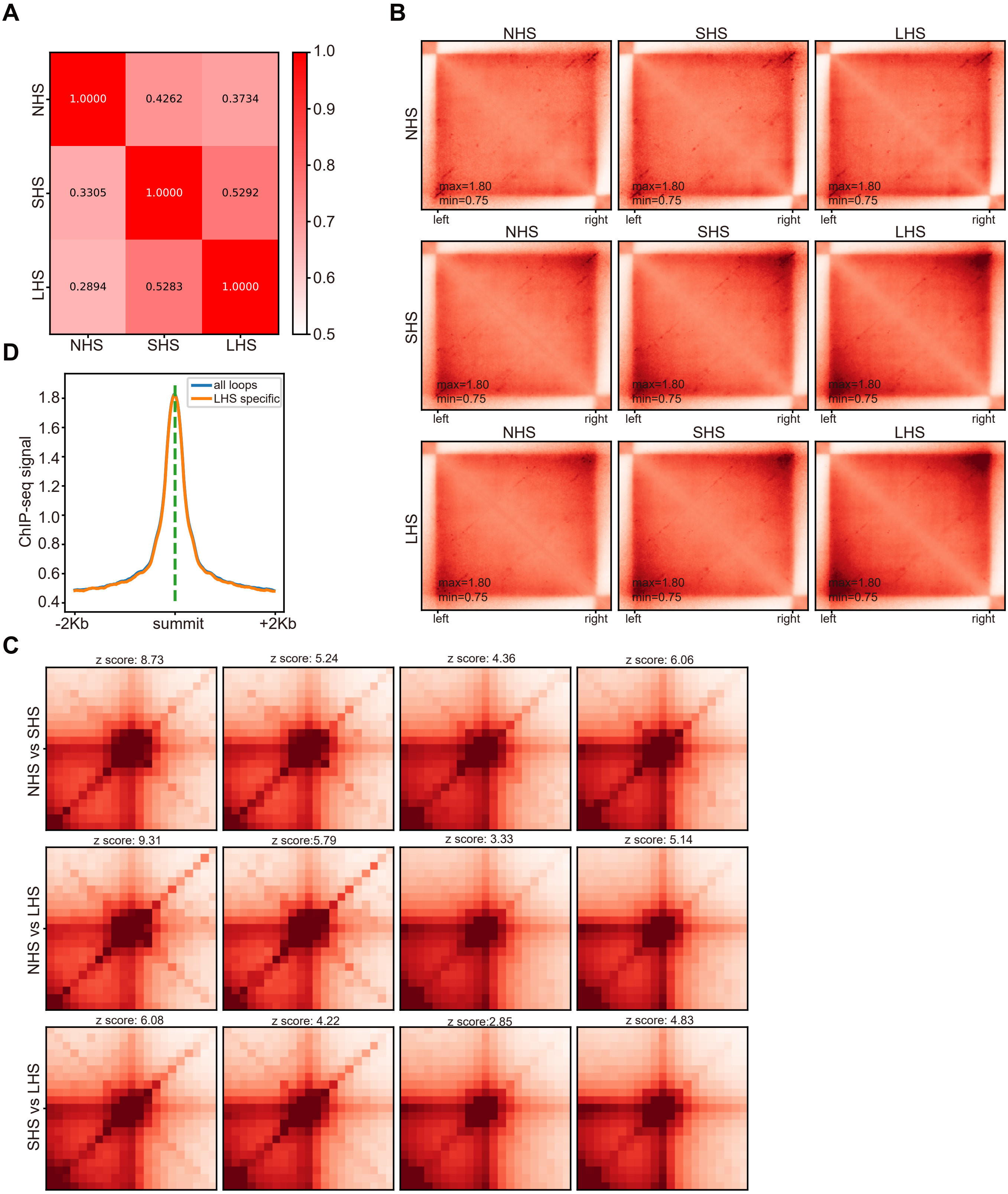

### Supplemental figure S8

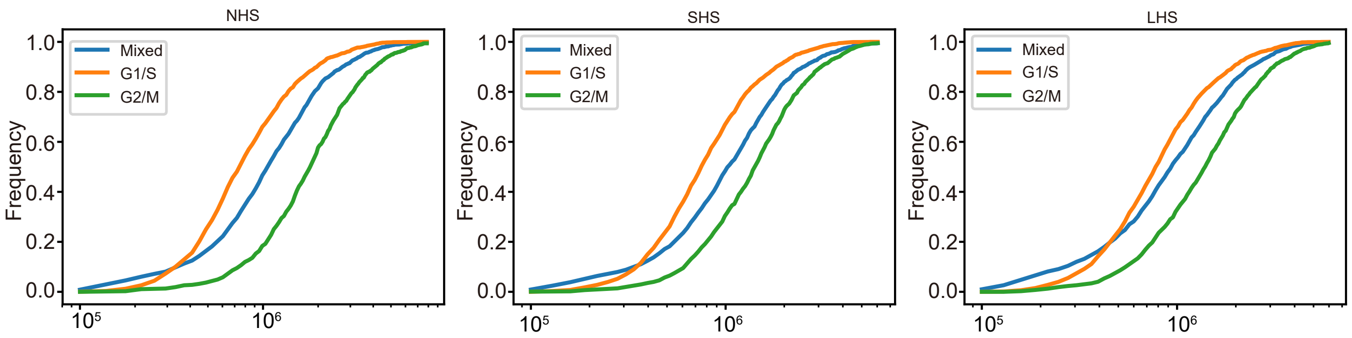
